# Synaptic vesicle fusion along axons is driven by myelination and subsequently accelerates sheath growth in an activity-regulated manner

**DOI:** 10.1101/2020.08.28.271593

**Authors:** Rafael G Almeida, Jill M Williamson, Megan E Madden, Jason J Early, Matthew G Voas, William S Talbot, Isaac H Bianco, David A Lyons

## Abstract

To study activity-regulated myelination, we imaged synaptic vesicle fusion along single axons in living zebrafish, and found, surprisingly, that axonal synaptic vesicle fusion is driven by myelination. This myelin-induced axonal vesicle fusion was enriched along the unmyelinated domains into which newly-formed sheaths grew, and was promoted by neuronal activity, which in turn accelerated sheath growth. Our results indicate that neuronal activity consolidates sheath growth along axons already selected for myelination.

## Main

Ensheathment of axons by myelin greatly regulates their conduction properties, enabling fast saltatory conduction (1) and providing metabolic support to the axon (2–4). In vivo, myelin can be specifically targeted to individual axons in a complex milieu (5), and can be highly regulated throughout life (6). Indeed, dynamic changes to myelin sheaths have been proposed to adapt circuit function to novel experiences, including learning (7, 8), social interactions (9–11), as well as memory formation and recall (12–14). Neuronal activity is now known to regulate oligodendrocyte lineage progression (15). It is also thought that activity might directly regulate myelination in vivo, through the fusion of synaptic vesicles along specific axons, a premise supported principally by experiments that abrogate vesicular fusion (16–18). Nevertheless, how myelination along individual axons is dynamically regulated in intact neural circuits remains unclear, partly due to the difficulty in observing changes to myelin along individual axons over time in vivo. It has been suggested that the synaptic connections that oligodendrocyte progenitor cells (OPCs) can make with axons might transform into myelin sheaths in an activity-regulated manner (19, 20), but recent studies have indicated that the OPCs that are synaptically connected to axons are not necessarily those that generate myelinating oligodendrocytes (21). It also remains unclear whether the fusion of synaptic vesicles occurs directly along the axon onto myelin sheaths, or only at synaptic termini, where it would regulate myelination indirectly, for example by spillover onto nearby axonal regions, or by driving changes in parameters such as axonal caliber that could influence myelination. Furthermore, it is not known whether more active axons exhibit greater levels of synaptic vesicle fusion that bias the initial selection of those axons for myelination, or whether highly active axons only release the contents of synaptic vesicles after axonal selection and sheath formation to consolidate myelination.

Here, we aimed to elucidate the mechanisms by which synaptic vesicle fusion and neuronal activity regulate the myelination of single axons over time in a living intact neural circuit. To address how vesicular fusion relates to and regulates myelination, we first aimed to define where and when synaptic vesicles fuse along individual axons in relation to their myelin sheaths. To do so, we performed live-cell imaging in developing zebrafish, which enables optical and genetic access to single neurons in an intact nervous system over time. We first characterized synaptic vesicular fusion along individual reticulospinal axons in the spinal cord, which become progressively myelinated in an activity/ synaptic vesicle fusion-dependent manner from 3 days post-fertilization (dpf) onwards (17). To do so, we used SypHy (22), in which synaptophysin, a transmembrane protein specifically localized to synaptic vesicles (23–25) is fused to four pH-sensitive pHluorin molecules. The pHluorin molecules face the acidic vesicle lumen, where their fluorescence is quenched. SypHy only fluoresces when the pH is neutralized upon synaptic vesicle exocytosis (Fig. 1a). The high signal-to-noise ratio provided by the multiple pHluorin molecules of this SypHy reporter enabled in vivo imaging of vesicular fusion. As previously reported (26, 27), a fraction of synaptophysin molecules reside on the plasma membrane, reflected by residual SypHy fluorescence that revealed individual reticulospinal axon morphology (Fig. 1b). Reticulospinal neurons have a large principal axonal shaft that is myelinated over time and which projects from the brain through the spinal cord, as well as regularly-spaced collateral branches that remain unmyelinated and contain presynaptic terminals (28). 1 Hz imaging of reticulospinal neurons at 4-5dpf, during active myelination, revealed bright, focal fluorescence increases at collaterals with presynaptic terminals, as expected, and also along the axon shaft (hereafter designated ‘axonal’ events) (Fig. 1c-d). Although SypHy event frequency was variable between individual neurons, axonal and collateral SypHy events were, on average, of similar frequencies (Fig. 1e), amplitudes and durations (Fig. S1a-b, d-e). Both axonal and collateral SypHy activity were blocked by co-expression of mCherry-tagged botulinum toxin B (BoNT-B) (29) (Fig. 1e, S1c-e) in individual neurons, which cleaves synaptobrevin, a vesicular transmembrane protein essential for exocytosis, validating events at both locations as bona-fide vesicular fusion (30). Most SypHy events (>75%) along control axons occurred in discrete locations with no subsequent displacement of fluorescent puncta and are thus likely to represent sites of focal vesicular fusion. A minority showed some displacement and also appeared insensitive to BoNTB (Fig. S1f-g), and so were not considered further. Collectively, our data reveal that in vivo, synaptic vesicles fuse not only at presynaptic terminals, as expected, but also along the length of reticulospinal axons at stages during which they become myelinated and where no presynaptic specializations have been reported. This suggests that synaptic vesicles could release their contents directly onto myelinating processes and/ or established sheaths, potentially regulating the formation, growth or maintenance of myelin.

**Fig 1.**
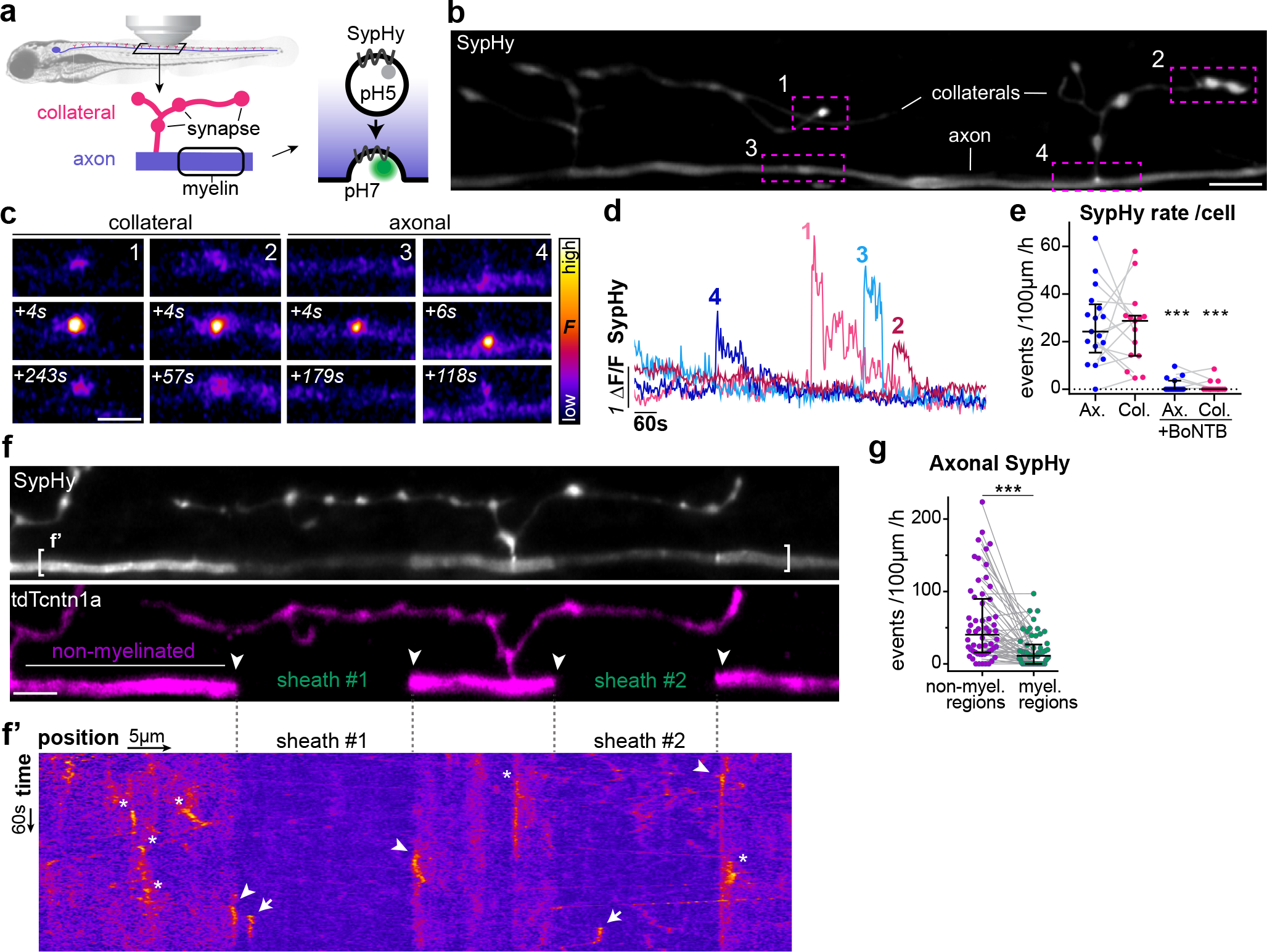
Axonal vesicular fusion is enriched in non-myelinated regions in vivo. a. Morphology of reticulospinal axons in the developing zebrafish spinal cord. SypHy consists of 4 molecules of pH-sensitive pHluorin fused to an intraluminal loop of rat synaptophysin (syp), a synaptic vesicle membrane protein. Exocytosis neutralises pH and dequenches SypHy signal. b. Individual SypHy+ reticulospinal axon (dorsal up) with synapse-bearing collateral branches. c-d. SypHy events at collaterals and axon and fluorescence time-courses. e. SypHy frequency is similar in axons and collaterals (each line shows respective events of same cell); and abolished in BoNTB+ neurons (p=0.82 Ax vs Col; p<0.001 Ax vs Ax+BoNTB; p<0.001 Col vs Col+BoNTB; Mann-Whitney test, 17 control axons from 16 animals; 15 BoNTB axons from 15 animals). f. SypHy and tdTcntn1a co-expression reveals subcellular distribution of axonal vesicular fusion, including in sheaths (flanked by arrowheads) and non-myelinated regions. Basal SypHy signal is decreased along sheaths, suggesting surface SypHy displacement by myelin. f’. SypHy kymograph of bracketed region in f, revealing vesicular fusion in non-myelinated regions (asterisks and arrowheads at sheath borders) and myelinated regions (arrows). g. Within myelinated axons, SypHy frequency is higher in non-myelinated regions (p<0.0001, Wilcoxon matched-pairs signed rank test, 58 myelinated axons from 53 animals). Scale bars: 5μm (b, f), 2μm (c). Graphs display median and interquartile range.

To directly examine the relationship between axonal vesicular fusion and the myelination of individual axons, we combined SypHy together with tdtomato-contactin1a (tdTc-ntn1a), our axonal reporter of myelination that is excluded from areas of the axon ensheathed by myelin (17, 31). We readily detected SypHy events along axons undergoing myelination (Fig. 1f-f’), with similar amplitudes and durations in myelinated (tdTcntn1a^-^) and non-myelinated (tdTcntn 1a^+^) regions (Fig. S2b), showing that they are equally detectable in both locations. Unexpectedly, however, we observed an over three-fold higher frequency of SypHy events in non-myelinated regions of axons undergoing myelination (Fig. 1g). This observation contradicts the premise that the principal location of synaptic vesicle fusion is directly under the newly forming or actively elongating myelin sheaths, and suggests instead that vesicular fusion occurs in regions into which sheaths are going to form and/ or into which they are about to grow.

To examine the relationship between axonal vesicular fusion and the onset of myelination in more detail, we analysed axonal SypHy activity in reticulospinal axons that were not yet myelinated (Fig. 2a-d). We found axonal SypHy activity was less frequent in axons not yet undergoing myelination than in axons undergoing myelination (Fig. 2e). In contrast, SypHy event frequency at collaterals, where presynaptic terminals are located, was comparable to that of myelinated axons (Fig. 2f). The observation that the onset of myelination coincides with a specific increase in axonal but not collateral vesicular fusion suggests that myelination itself might in fact stimulate vesicular fusion from axons. To test this possibility, we specifically reduced CNS myelin formation by genetically disrupting zebrafish *myrf*, which encodes an oligodendrocyte-specific transcription factor required for myelin production (Bujalka et al., 2013; Emery et al., 2009; Koenning et al., 2012). At 4-5dpf, most reticulospinal axons in wildtype siblings were in the process of being myelinated (median 30% axonal length myelinated, 25 axons); whereas myrf mutant siblings had little myelin (median of 0% axonal length myelinated, 12 axons) (Fig. 2g-i). We found that axonal SypHy events were specifically decreased in *myrf* mutants (Fig. 2j), whereas collateral SypHy event frequency was not significantly affected (Fig. 2k). These data indicate that, surprisingly, myelination itself drives vesicular fusion along myelinated axons, rather than vesicular fusion biasing the initial targeting and selection of specific axons for myelination. Given that myelination-induced axonal vesicular fusion becomes enriched in non-myelinated regions (Fig. 1g), how could it regulate myelination? Does it affect sheath growth after the myelinating process becomes targeted to the axon? To address precisely where and when axonal vesicular fusion occurs, we examined the subcellular distribution of SypHy events along single myelinated axons in more detail. In 70 myelinated axons from 57 animals, we first found that SypHy activity in non-myelinated regions correlated positively with the extent of an axon’s myelination (Fig. S3a), such that axons with more SypHy events in their non-myelinated regions were also the axons with more myelin coverage along their length. In contrast, SypHy activity in myelinated regions did not correlate with the extent of an axon’s myelination.

**Fig 2.**
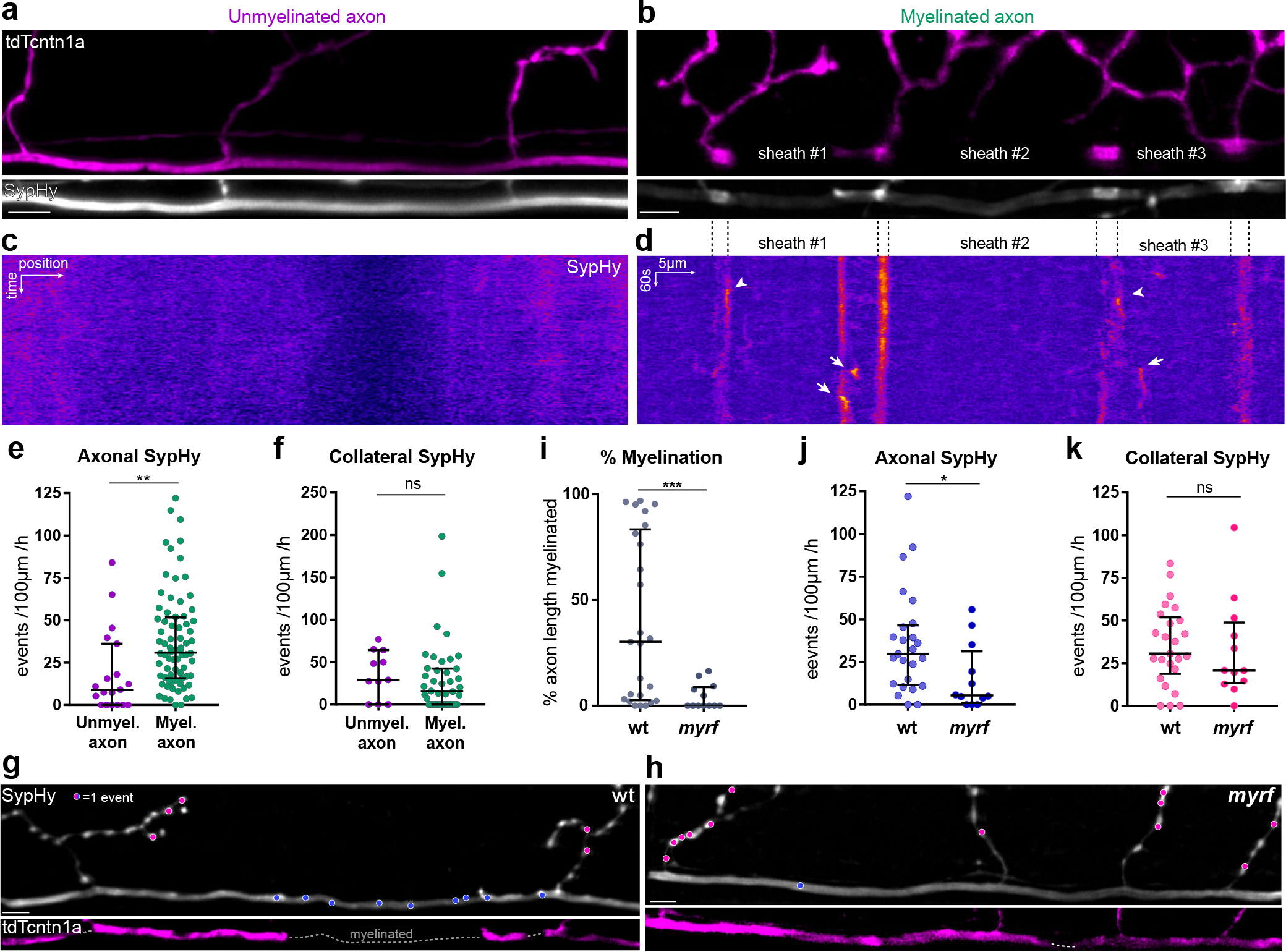
Myelination promotes axonal vesicular fusion. a-b. SypHy and tdTcntn1a profile of a not-yet myelinated (a) and a myelinated (b) reticulospinal axon. In more fully myelinated axons, tdTcntn1a becomes enriched at putative nodes of Ranvier between adjacent sheaths. c-d. Corresponding SypHy kymographs of bracketed segments in a-b. e-f. Axonal (e) but not collateral (f) SypHy frequency in myelinated axons is greater than in axons yet to be myelinated (e: p=0.001; f: p=0.30; Mann-Whitney test, e: 19 unmyelinated axons from 19 animals and 75 myelinated axons from 68 animals). g-h. SypHy activity (1 dot = 1 event) and myelination profile of axons in wildtype (g) and hypomyelinated myrf mutants (h). i. Myelin coverage in wt and myrf (p=0.002, Mann-Whitney test, 25 axons from 19 wt animals and 12 axons from 11 myrf animals). j-k. Axonal (j) but not collateral (k) SypHy frequency is decreased in myrf mutants (j: p=0.022; k: p=0.476; Mann-Whitney test, same N as i). Scale bars: 5μm (a-c, g-h), 60s (c-d). Graphs display median and interquartile range.

We next observed that within non-myelinated regions, many events were localised adjacent to the ends of individual myelin sheaths (within 3μm, Fig. 1f’, 3a-c, arrowheads), at a frequency higher than expected if they were uniformly distributed along the entire non-myelinated regions (Fig. 3d). In axons undergoing myelination, these regions immediately adjacent to growing sheaths - into which myelin sheaths are likely to grow - are called heminodes since they often display an enriched localisation of proteins that ultimately cluster at nodes of Ranvier as neighbouring sheaths grow towards one another (32–34). Co-expression of SypHy and nfasca-mCherry, a nodal cell adhesion molecule, showed that the non-myelinated regions adjacent to the ends of myelin sheaths were indeed enriched in nfasca during the myelination of zebrafish reticulospinal axons (Fig. S2c), confirming that they had heminodal characteristics.

**Fig 3.**
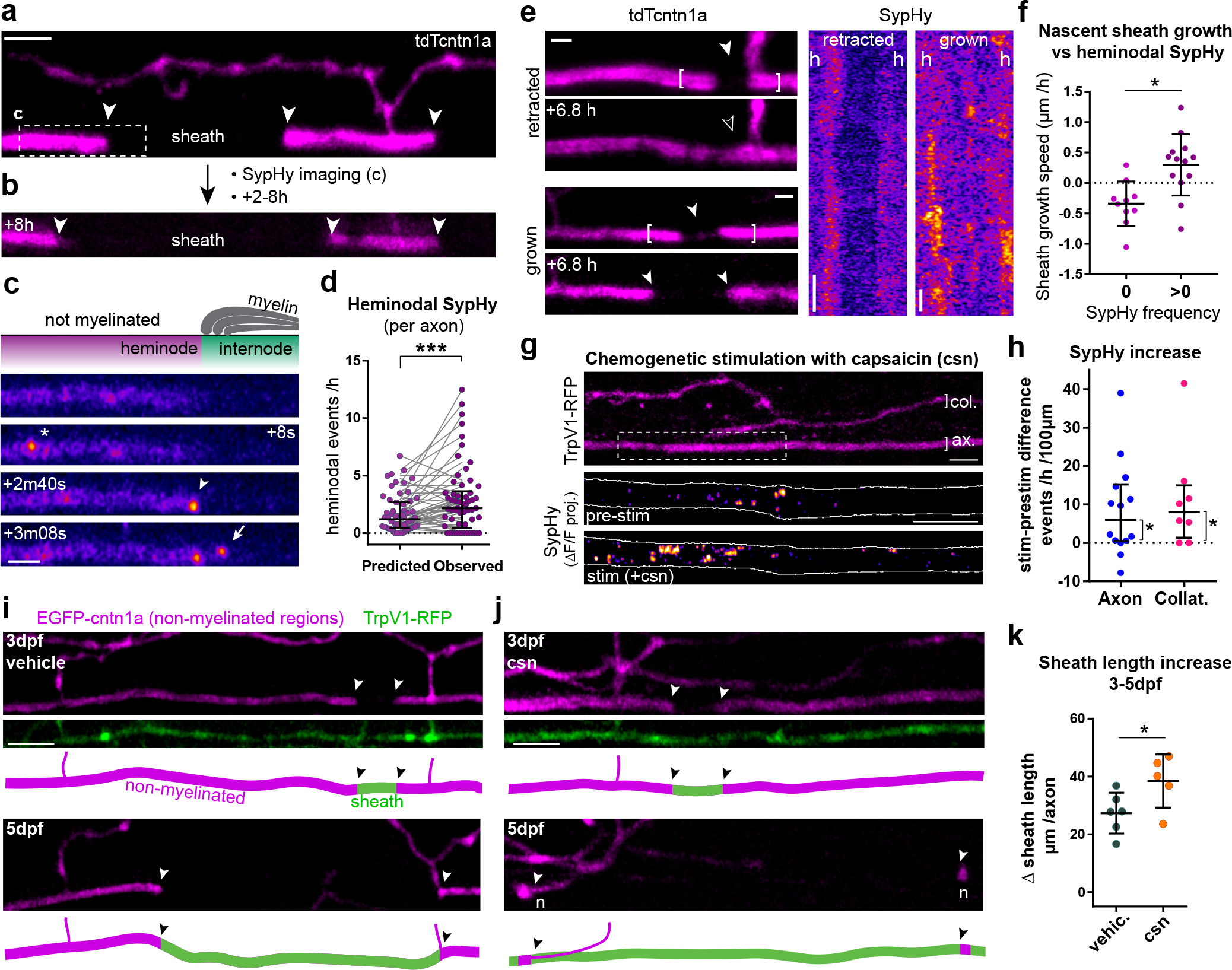
Axonal vesicular fusion promotes myelination. a-b. tdTcntn1a reveals myelin growth, here over 8h (note sheath growth, arrowheads). c. SypHy activity of box in a, with events along non-myelinated regions (asterisks), putative heminodes (arrowheads) and myelinated regions (arrows). Related to kymograph in Fig. 1f’. d. Observed heminodal Syphy frequency per axon is higher than predicted if non-myelinated events were uniformly distributed (p=0.0004, Wilcoxon matched-pairs signed rank test, 58 myelinated axons from 53 animals). e. SypHy activity in eetracting and growing nascent sheaths (h=heminode). All complete retractions were of sheaths <5μm. f. Nascent sheaths with heminodal SypHy activity grow faster (nascent: p=0.003, Student’s t-test, 23 nascent sheaths from 14 axons from 13 animals). g. SypHy in TrpV1 + axon before and during csn treatment, ΔF/F projection over 11min (before) and 12min (+csn). h. Capsaicin increased axonal and collateral SypHy in TrpV1+ axons above baseline (Difference from zero: p=0.013 axonal, p=0.031 collateral, Wilcoxon signed-rank test, 14 axons and 8 collaterals from 14 and 8 animals respectively). i-j. Single TrpV1 + axons treated with vehicle (i) or csn (j) throughout their myelination (arrowheads – putative heminodes; n - putative nodes of Ranvier). k. Stimulation promotes sheath growth (3-5dpf change: p=0.04, Student’s t-test). 6 control axons and 5 stimulated axons from 6 and 5 animals respectively. Scale bars: 2μm (c, e), 5μm (a, g, i-j), 60s (e). Graphs in f and k display mean and standard deviation, d and h show median and interquartile range.

The enrichment of SypHy events at sites into which sheaths could grow suggested that localised vesicular fusion might promote sheath elongation into those regions. In agreement with this premise, we found that in 162 sheaths imaged along 61 axons (in 54 animals), sheath length correlated positively with SypHy event frequency in the heminodal regions. We saw no correlation between sheath length and sypHy activity under the sheath itself (Fig. S3b). To relate axonal vesicular fusion and the growth rate of individual sheaths, we imaged the myelination profiles of 39 individual axons before and 2-8h after SypHy imaging (Fig. 3a-c). Overall, heminodal SypHy event frequency correlated positively with the speed of sheath growth (Fig. S3c). 71/86 sheaths elongated over this time, while only 14/86 shrank, half of which retracted completely. Interestingly, nascent sheaths <5μm were equally likely to grow (48%) or shrink (52%) while sheaths >5μm rarely shrank (10%) (Fig. S3d). This length-dependent fate suggests that newly-formed sheaths are vulnerable to retraction but that factors promoting their elongation past a 5μm threshold ensure their stable formation. Indeed, all complete retractions were of sheaths <5μm. Remarkably, 85% of nascent sheaths with heminodal SypHy activity elongated, compared to only 20% of those with zero SypHy activity (Fig. 3e-f). Thus, vesicular fusion adjacent to sites of sheath formation positively correlates with faster sheath growth, suggesting that activity is one of the factors that can drive myelin elongation past a threshold that promotes their maintenance. We previously showed that constitutive abrogation of vesicular fusion in individual reticulospinal axons reduced myelin sheath number and length (17). To test whether the activity of individual axons can also enhance the growth of myelin sheaths, we sought to enhance vesicular fusion along single reticulospinal axons. We expressed tagRFP-tagged rat TRPV1, a cation channel specifically activated by capsaicin, which drives neuronal excitation and activity-dependent vesicular fusion, and has been used in zebrafish to increase the firing frequency of spinal neurons (Chen et al., 2016). 1μM capsaicin (but not vehicle) increased reticulospinal intra-axonal calcium activity (as assessed with axon-tethered GCaMP7s (35)) in individual TRPV1-RFP+ axons compared to their baseline, and to control axons (Fig. S3e-h), indicating a specific increase in neural activity in treated transgenic axons with our approach. Capsaicin also significantly increased the frequency of both axonal and synaptic SypHy events (Fig. 3g-h), with 10/12 TrpV1+ axons showing increased axonal SypHy (overall average 1.6 fold-increase). These data indicate that SypHy-reported axonal vesicular fusion is activity-regulated and is increased with our chemogenetic approach. We next increased vesicular fusion in individual axons throughout their period of myelination by treating larva with capsaicin every day for 4 hours from 3-5dpf. We imaged their myelination profile using EGFPcntn1a (Fig. 3i-j) and found that sheaths on stimulated axons grew significantly more, from 8±4μm at 3dpf to 45±6μm by 5dpf in stimulated axons, compared to 36±5μm in controls (Fig. 3k). Thus, stimulating activity specifically promotes an increase in myelin sheath growth. Since activity-regulated vesicular fusion along axons is itself promoted by the onset of myelination (Fig. 2), our data reveal that myelination itself triggers the activity-regulated mechanism that promote further continued myelination of specific individual axons in developing neural circuits. We and others have previously shown that vesicular fusion is required for appropriate myelination along axons (16, 17). Here, we make the unexpected observation that the onset of myelination itself stimulates localised axonal vesicular fusion, which in turn promotes sheath growth. This contrasts with the proposal that vesicular fusion, from axon-OPC synapses or otherwise, provides the signal that attracts myelinating processes to more active axons (5, 36, 37). Our data instead suggest a model where oligodendrocytes select specific axons in an activity-independent manner; and this sheath formation then promotes axonal vesicular fusion, which consolidates sheath formation and elongation, consistent with previous observations that impaired vesicular release reduced, but was not absolutely required for myelination along single axons (17, 18). The molecular mechanisms by which myelination promotes vesicular release and by which vesicular release in turn promotes myelination remain to be defined. For instance, does myelination locally concentrate vesicular fusion machinery in the underlying axon, or drive other localised changes, e.g. to axonal diameter, that facilitate vesicular trafficking or fusion (38, 39)? Do similar mechanisms regulate the fusion of synaptic vesicles and other vesicles, e.g. those containing nodal components (33, 40), along axons? It will also be important to test how myelination affects vesicular fusion along different types of axons, given that vesicular fusion has been shown to affect the myelination of only some neuronal subtypes, and not others (17).

Our data clarify how activity-regulated vesicular fusion in turn affects myelination. We and others have previously shown that abrogation of vesicular release from neurons reduces the number and length of myelin sheaths (16, 17). However, it has remained unclear to what extent sheath formation and elongation along axons are distinctly regulated by activity. Here, we propose that activity-regulated vesicular fusion promotes the stable formation of sheaths by promoting their growth and elongation. The temporal resolution afforded to us by our model enabled us to determine that once sheaths reach a length of around 5μm they only ever elongate, whereas shorter sheaths are prone to shrinking and retraction. We show that axonal vesicular fusion promotes nascent sheath growth and thus enables their stable formation. As axons become fully myelinated, activity/vesicular fusion likely regulate sheath thickness through promoting their growth (41, 42), but further analysis in mature circuits will determine how long vesicular fusion-regulated myelination remains operational. In addition, the cargo of synaptic vesicles promoting myelin elongation remains to be identified. Reticulospinal axons are glutamatergic (43, 44), and vesicular glutamate release has previously been implicated in myelination (20, 45). Defining when and where this occurs, and which receptors on myelinating processes mediate the effects of vesicular cargos, as well as the functional consequences of activity-regulated myelination of single axons in vivo remain important questions to address in the future.

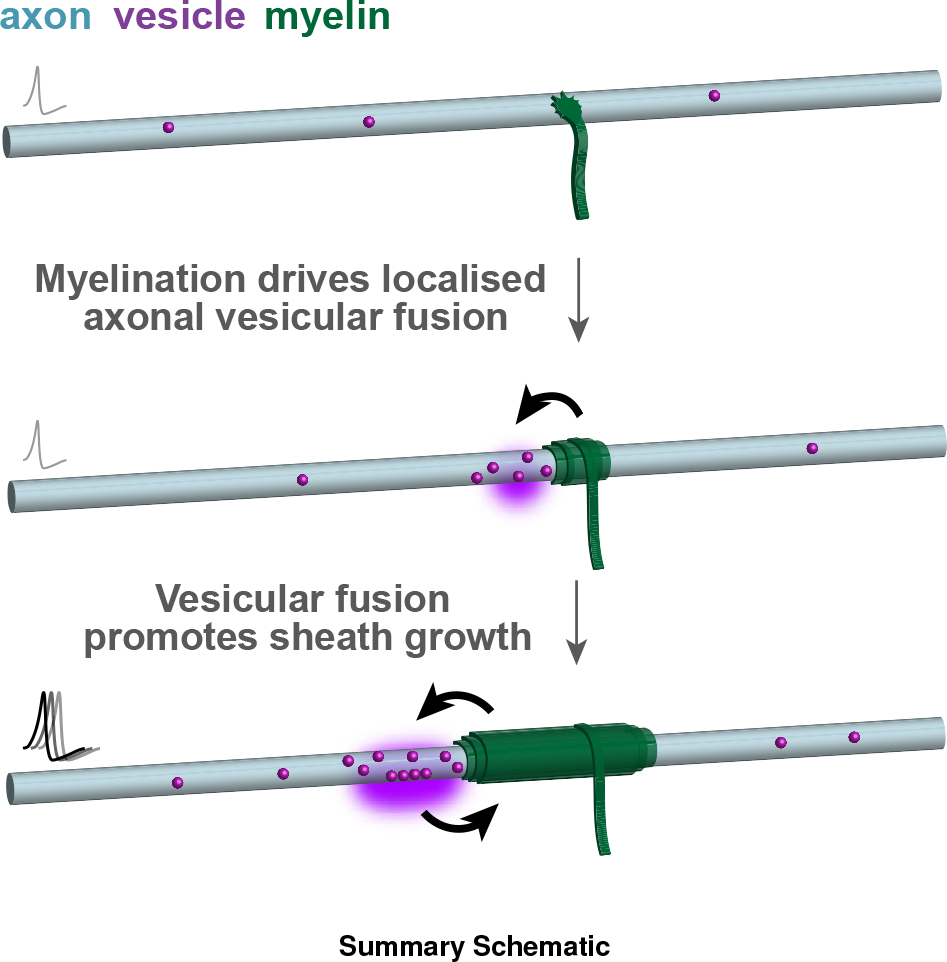

In summary, our in vivo imaging of synaptic vesicle fusion from axons undergoing myelination has revealed an unexpected model whereby the onset of myelination induces localised activity-regulated vesicular fusion along the axon that in turn promotes myelin growth (see Summary Schematic). This represents an efficient mechanism to ensure the timely and robust myelination of specific CNS axons and circuits.

## Acknowledgements

We thank members of the Lyons lab, Peter Brophy, Tim Czopka, Michael Cousin, Jonah Chan, Matthew Livesey, Dies Meijer, and Matthew Swire for helpful comments and critique of our manuscript. UAS:BoNTB-mCherry plasmid was a gift from Claire Wyart; islet1:GAL4VP16,4xUAS:TRPV1-RFPT plasmid was a gift from David Prober; pcDNA3-SypHluorin 4x (S4x) was a gift from Stephen Heinemann & Yongling Zhu (Addgene plasmid 37005); pGP-CMV-jGCaMP7s was a gift from Douglas Kim & GENIE Project (Addgene plasmid #104463). This work was supported by a Medical Research Council (MRC) project grant (MR/P006272/1) and Wellcome Trust Senior Research Fellowships (102836/Z/13/Z and 214244/Z/18/Z) to D.A. Lyons, a University of Edinburgh Ph.D. Tissue Repair Studentship Award (MRC Doctoral Training Partnership MR/K501293/1) and the Wellcome Trust Four-Year Ph.D. Program in Tissue Repair (Grant 108906/Z/15/Z) to JMW, a Wellcome Trust Edinburgh Clinical Academic Track PhD studentship to MEM, a Sir Henry Dale Fellowship from the Royal Society & Wellcome Trust (101195/Z/13/Z) and a UCL Excellence Fellowship to IHB.

## Methods

### Zebrafish Lines and Maintenance

All zebrafish were maintained under standard conditions (46, 47) in the Queen’s Medical Research Institute BVS Aquatics facility at the University of Edinburgh. Studies were carried out with approval from the UK Home Office and according to its regulations, under project licenses 60/8436, 70/8436 and PP5258250. Adult animals were kept in a 14 hours light and 10 hours dark cycle. Embryos were kept at 28.5°C in 10mM HEPES-buffered E3 Embryo medium or conditioned aquarium water with methylene blue. Embryos were staged according to (48), and analyzed between 2-7dpf, before the onset of sexual differentiation. The following existing transgenic line was used to drive expression in reticulospinal neurons: Tg(KalTA4u508)u508Tg (49). The existing panneuronal driver line Tg(huC:Gal4) was also used (Mensch et al., 2015). The following lines were generated in this study: *myrf*^UE70^ (described below); Tg(10xUAS:Sy4xpHy; cryaa:mCherry); Tg(10xUAS:GCaMP7s; cryaa:mCherry); Tg(10xUAS:EGFP-cntn1a); Tg(10xUAS:TrpV1-tagRFPt). Throughout the text and figures, ‘Tg’ denotes a stable, germline-inserted transgenic line; and ‘SypHy’ denotes the variant containing four pHluorin molecules, also called ‘Sy4xpHy’ (22).

### *myrf*^UE70^ generation, genotyping and analysis

The *myrf*^UE70^ allele was generated by injection of Cas9 mRNA and a sgRNA targeting the second exon of *myrf* (target sequence: CATTGACACCAGTATCCTGG) into fertilized wildtype embryos at one-cell stage. Potential founders were grown to adulthood and the UE70 allele isolated from their offspring. UE70 consists of an indel (ΔCC+A) that disrupts the reading frame and introduces a premature stop codon. Homozygous *myrf*^UE70^ mutants have reduced numbers of oligodendrocytes and exhibit myelination of the spinal cord; an in-depth description of this mutant line will be published elsewhere (Madden et al., in prep).

For SypHy analysis in *myrf*^UE70^ mutants, *myrf*^UE70^ heterozygous parents carrying the Tg(KalTA4u508) neuronal driver were in-crossed, and offspring were injected at one-cell stage with SypHy-UAS-tdTomato-cntn1a and tol2 mRNA, as detailed below. Following SypHy imaging at 4-5dpf, individual larvae were genotyped using primers Myrf F and Myrf R (Table S1) followed by digestion of the PCR product with restriction enzyme PspGI, which cleaves the wildtype PCR product into 131bp and 157bp-long fragments, but not the mutant product, since the *myrf*^UE70^ allele consists of a frameshifting indel which abolishes the PspGI site. Subsequent analysis was performed blinded to genotype.

### Generation of Tol2kit-compatible entry vectors

To generate tol2kit (50) compatible entry vectors, we amplified the relevant coding sequences by PCR with Phusion and primers as indicated in Tables S1 and S2 and recombined 25fmol of purified PCR product with 75ng of pDONR221, pDONRP4P1R or pDONRP2RP3, as appropriate, in a BP reaction using BP Clonase II. Additional/alternative construction details are as follows. p3E-(zf)tdTomato-cntn1a: we replaced the tagRFPt coding sequence in p3E-tagRFPt-cntn1a (31) with a zebrafish codon-optimized non-repetitive sequence of tdTomato. We amplified zf-tdTomato from a gBlock (IDT DNA, full sequence available upon request) with primers *zftdTomato fwd* and *zftdTomato rev* and the p3E-tagRFPtcntn1a backbone (excluding tagRFPt) with primers *zfcntn1a signal rev* and *zfcntn1a Fwd Phos*, and ligated both using T4 DNA ligase. pME-axonGCaMP7s: we digested pME-jGCaMP7s at the start codon with NcoI (GCCA*<u>CCATGG</u>*, NcoI sequence underlined) into which we ligated two annealed primers, GAP43-fwd and GAP43-rev, encoding the di-palmitoylation motif of murine GAP-43 flanked by overhanging NcoI-compatible ends. This motif directs and enriches jGCaMP7s localization to axons compared to non-specific membrane tethers (35). p3E-nfasca-mCherry: zebrafish *nfasca* transcripts were amplified by RT-PCR from total brain RNA using primers EcoRI-Kozac-nfasca-F and nfasca-NotI-R (based on cDNA clone IMAGE: 3817913, accession CU458816), cloned into pCRII-TOPO and sequenced. A zebrafish isoform that contains a mucin-like domain (accession FJ669144) was considered as the neuronal form of *nfasca*, similar to the mammalian neuronal isoform NF186. This zebrafish NF186-like cDNA cloned into expression plasmid pUAS-NF186. The HA tag (YPYDVPDYA) was inserted between amino acids 35 and 36 following a predicted secretion signal sequence (SignalP) by recombinant PCR using equimolar amounts of a PCR product containing *nfasca* coding nucleotides 1-105 and the HA tag (amplified with primers EcoRI-Kozac-nfasca-F and nfasca-HA-R) and a PCR product containing nucleotides 106-767 (amplified with primers nfasca-HA-F and nfasca-AgeI-R), and primers EcoRI-Kozac-nfasca-F and nfasca-AgeI-R. The resulting PCR product was cloned into pCRII-TOPO and then used to replace the 5’ end of NF186 in pUAS-NF186 using an EcoRI site upstream of the start codon and an AgeI site downstream of the start codon. The resulting full-length HA-NF-186 fusion was then cloned into a modified version of the Tol2 transgenesis vector pBH-UAS (Michael Nonet, University of Washington) yielding pBH-UAS-HA-NF186. The mCherry coding sequence was then PCR-amplified with primers SacI-nfasca-mCherry-F and mCherry-NotI-R, cloned into pCRII-TOPO, and then used to join the HA-NF186 fusion with the mCherry fusion by ligating a BsiWI/SacI NF186-like cDNA fragment, a SacI/NotI mCherry fragment, and a BsIWI/NotI pBH-UAS-HA-NF186 fragment. This yielded the expression plasmid pBH-UAS-HA-NF186-mCherry, from which the HA-NF186-mCherry coding sequence was amplified to make p3E-nfasca-mCherry using the primers indicated in Table S2. Coding sequences in all 5’-entry vectors are in the reverse orientation and contain a polyadenylation signal, to be compatible with a ‘Janus’ configuration (51) following LR recombination with a middle-entry vector containing (palindromic) UAS and a 3’-entry vector containing another coding sequence in the forward orientation. The sequences of all entry vectors were verified by Sanger sequencing.

### Generation of Tol2 expression/transgenesis constructs

To generate final Tol2 expression vectors, 10fmol of each entry vector as indicated in Table S3 and 20fmol of destination vector pDestTol2pA2 form the tol2kit or pDestTol2pA2-cryaa:mCherry(Berger and Currie, 2013) (Addgene #64023) were recombined in a LR reaction using LR Clonase II Plus. 3-4 clones were tested for correct recombination by digestion with restriction enzymes.

### Generation of transgenic lines

Transgenic lines were generated by injecting 5-10pg of the appropriate plasmid DNA with 25-50pg *tol2* transposase mRNA into wild-type zebrafish eggs at the one-cell stage to promote transgenesis. The plasmid used to make the Tg(UAS:EGFP-cntn1a) has been previously described (17). Founder animals were identified by screening F1 offspring for transgenesis markers, and F1 offspring were raised to generate stable transgenic lines. Lines were not screened for single transgene insertions.

### Microinjection/Single cell labelling

One-cell stage Tg(HuC:Gal4) or Tg(KalTA4u508) eggs were injected with 1-5pg of plasmid DNA containing UAS-driven SypHy or other reporters as indicated with 25pg of *tol2* transposase mRNA, to mosaically express axonal reporters in RS neurons. one-cell stage Tg(KalTA4u508; UAS:TRPV1-RFP) eggs were injected with 5pg 10xUAS:axon-jGCaMP7 with 25pg tol2 transposase mRNA to mosaically label RS neurons with axon-localised GCaMP7.

### Live-imaging

For SypHy imaging, larva expressing SypHy in reticulospinal axons were obtained either by crossing Tg(KalTA4u508) with Tg(UAS:SypHy) fish or by injecting UAS:SypHy plasmid and *tol2* mRNA into Tg(KalTA4u508) fertilized eggs. At 3-5dpf, larvae were paralysed by a 5minute bath-application of 1.5mg/ml mivacurium chloride (Abcam) in E3 embryo medium and immobilised in 1.5% low melting-point agarose on their sides. Animals were imaged using a Zeiss LSM880 confocal with Airyscan in Fast mode and a Zeiss W Plan-Apochromat 20x/1.0 NA water-dipping objective, and a 488nm (SypHy), 568nm (tdTomato, tagRFPt) and 594nm (mCherry) laser. We used 3-4.5X zoom and acquired 1500-2300 (X) x 50-500 (Y) pixels, sampling 100-140μm of axonal length (16 pixels/μm), over a small z-stack (4-15 z-slices, 1-5μm z-step) that sampled the depth of the axon and collateral branches. This was acquired repeatedly at a frequency of 0.6-2.3 Hz (mean 1.1Hz), for 5-20 minutes. For experiments in which axons co-expressed a static red fluorescent marker (obtained by injecting SypHy-UAS-BoNTBmCherry, SypHy-UAS-tdTomatocntn1a or SypHy-UAS-TrpV1RFP into fertilized Tg(KalTA4u508) eggs), the red channel was acquired once, in an optimally-sectioned z-stack, in the same region as SypHy, with higher pixel dwell time to reduce noise. For imaging SypHy with a coexpressed dynamic red marker (obtained by injecting SypHy-UAS-nfascamCherry into fertilized Tg(KalTA4u508) eggs), we acquired every line of each channel sequentially to maximize temporal resolution. For axonal Ca^2+^ imaging (of larvae obtained by injecting axon-GCaMP7s-UAS-TrpV1RFP into fertilized Tg(KalTA4u508) eggs), a higher resolution image of TRPV1-RFP and axon-GCaMP7 expression was acquired, then a 3-5 minute timelapse movie of GCaMP7 alone was acquired at 1Hz before and 15 minutes after vehicle or capsaicin treatment (detailed below).

### Chemogenetic stimulation

Capsaicin (Sigma-Aldrich) was prepared as a 5mM primary stock in 100% DMSO and stored at −80°C. For chronic treatments, the transgenic lines Tg(UAS:TRPV1-RFP) and Tg(UAS:GFP-Cntn1a) were crossed into the Tg(KalTA4u508) background to drive mosaic expression of both transgenes in reticulospinal neurons. Animals with individual reticulospinal axons expressing both transgenes were live-imaged at 3dpf as above and then returned to E3 embryo medium. Animals were treated with either vehicle (1% DMSO) or capsaicin (1μM capsaicin in 1% DMSO) in E3 embryo medium for 4 hours after imaging at 3dpf, and then every day until 7dpf. Animals were re-imaged as above at 5dpf to obtain time-course data, and were kept individually in 12-well plates in-between imaging sessions. For acute capsaicin treatment of axon-GCaMP7s and SypHy expressing larva, before imaging as detailed above, a small window of agarose was removed along the trunk of the animal to facilitate diffusion of 1μM capsaicin-containing (or vehicle) E3 embryo medium, which was allowed to equilibrate for 10 minutes before re-imaging.

### Image processing and analysis

We used Fiji/ImageJ and Python scripts for most image processing and analysis, and Adobe Illustrator for figure panels, using average or maximum-intensity projections and cropped representative x-y areas. SypHy time-lapses were pre-processed by bleach-correction with exponential curve fitting, and registration, where needed, using the template-matching plugin or custom written ImageJ macros that apply rigid transformation. Putative SypHy events were identified in the pre-processed (but otherwise raw) movies, aligned to a ΔF/F_avg_ movie (proportional increase over the all-time average intensity) to aid discrimination of the event start. For each potential event, we defined its region of interest using Fiji’s Wand tool to select the maximum intensity pixel and connected region that was over a third of the maximum fluorescence intensity. We used this ROI to determine the increase in fluorescence intensity relative to the baseline, defined as the average intensity in the ten frames preceding the event (F_0_), and only considered events with an increase that was 5-fold greater than the baseline standard deviation, lasting at least 5 frames. The amplitude was defined as the highest proportional increase that the ROI reached over the baseline (ΔF/F_0_) during the peak period, defined as the first ten frames. Duration was defined as the time until fluorescence decreased to within one standard deviation of the baseline. We also calculated the cumulative displacement of the event using the maximum intensity pixel and, where substantial (> 2μm) classified this displacement as unidirectional or bidirectional for axonal events, and anterograde or retrograde (away or towards the axon, respectively) for collateral events. We excluded from further analysis events that showed motion throughout their duration or >10μm displacement, but included those events that showed some, limited displacement (<10μm), and that were static for longer periods than they were moving (these could represent, for example, instances of kiss-and-run exocytosis). Collection of these parameters was automated using custom written ImageJ macros, but all individual events were manually inspected and parameters corrected where needed. SypHy event frequency was normalized to axonal or collateral length, which were measured in an average-intensity projection of all time frames; and to time imaged. Kymographs were made using the Fiji Multi Kymograph plugin.

For analysis of tdTomato-cntn1a expressing axons, myelin sheath location and lengths were inferred from tdTomato-cntn1a negative gaps that bridged the axon’s thickness, as before (17). The percentage of myelination was calculated as the summed length of tdTomato-negative gaps in the axon, including those that are incomplete at the edges of the field of view, divided by the total length of axon sampled in the field of view. We considered axons yet-to-be-myelinated if they had <5% myelination, and axons actively undergoing myelination if they had >5% myelination or >2 putative heminodes. Although we did not always sample axons at the same somite level, all images were taken at the mid-trunk level, approximately between somites 10-20. We classified the location of all axonal SypHy events as tdTomato+ or tdTomato-, and further classified tdTomato+ Syphy events as ‘heminodal’ when occurring at putative heminodal locations, defined as the first 3μm of tdTomato-positive axon bordering a tdTomato-negative gap. Collateral branching points were defined as a 3μm segment of axon centered on the collateral. Collaterals branching points were always located in non-myelinated tdTomato+ parts of the axon. We normalized SypHy frequencies to tdTomato+ length and tdTomato-length where appropriate. To calculate the predicted heminodal SypHy frequency in each axon, we scaled the overall SypHy frequency in the tdTomato+ part of the axon to the frequency in a 3μm window, expressed as the #events/3μm/h, which would reflect a uniform distribution, and compared this to the observed overall heminodal frequency divided by the number of heminodes. For sheath-centric analyses, we only considered complete gaps, i.e. excluding incomplete gaps at the edge of the field of view. We considered an heminode as a node if its length was <1μm and immediately flanked by another gap. For myelin growth analysis, we identified individual sheaths over time based on axonal landmarks, including collateral branches, which have unique morphologies. We considered sheaths ‘free’ to grow if they were not bordered by nodes or collaterals.

For axon-GCaMP7s Ca^2+^ analysis, a well-defined collateral branching from the main axon was defined as a ROI and background-subtracted proportional change over the average was calculated ΔF/F_avg_. The pre-treatment movie was considered a measure of baseline neuronal activity, and for the post-treatment movie, the proportional change in fluorescence ΔF/F_0_ was calculated using the average of pretreatment movie as F_0_. For analysis of individual axon myelination during chronic stimulation, one axon was analysed per animal, and sheath length inferred from FP-cntn1a gaps measured using Fiji as before (17).

### Quantification and statistical analysis

All graphs and statistical tests were carried out using GraphPad Prism. All data were averaged per biological replicate (N represents number of animals), except where otherwise noted (where N represents the number of different neurons or sheaths). Data were tested for normal distribution using D’Agostino & Pearson and Shapiro-Wilk normality tests. Normally distributed groups were compared using two-tailed unpaired Student’s t-test and non-normally distributed groups were compared using the Mann-Whitney U test or Wilcoxon matched-pairs signed rank test for paired data. For correlation analysis, we used both parametric Pearson’s test for linear correlation and non-parametric Spearman’s test for monotonic correlation, as indicated in Figure legends. We considering a difference significant when p<0.05, and indicate p values in figures as follows: no indication or ‘ns’ p>0.05, ‘*’ p<0.05, ‘**’ p<0.01, ‘***’ p<0.001. Error bars illustrate mean ± standard deviation for normally distributed data or median and interquartile range for non-normally distributed data, and further details on statistical tests are indicated in Figure legends.

## Supplementary Tables

**Table S1 -.**
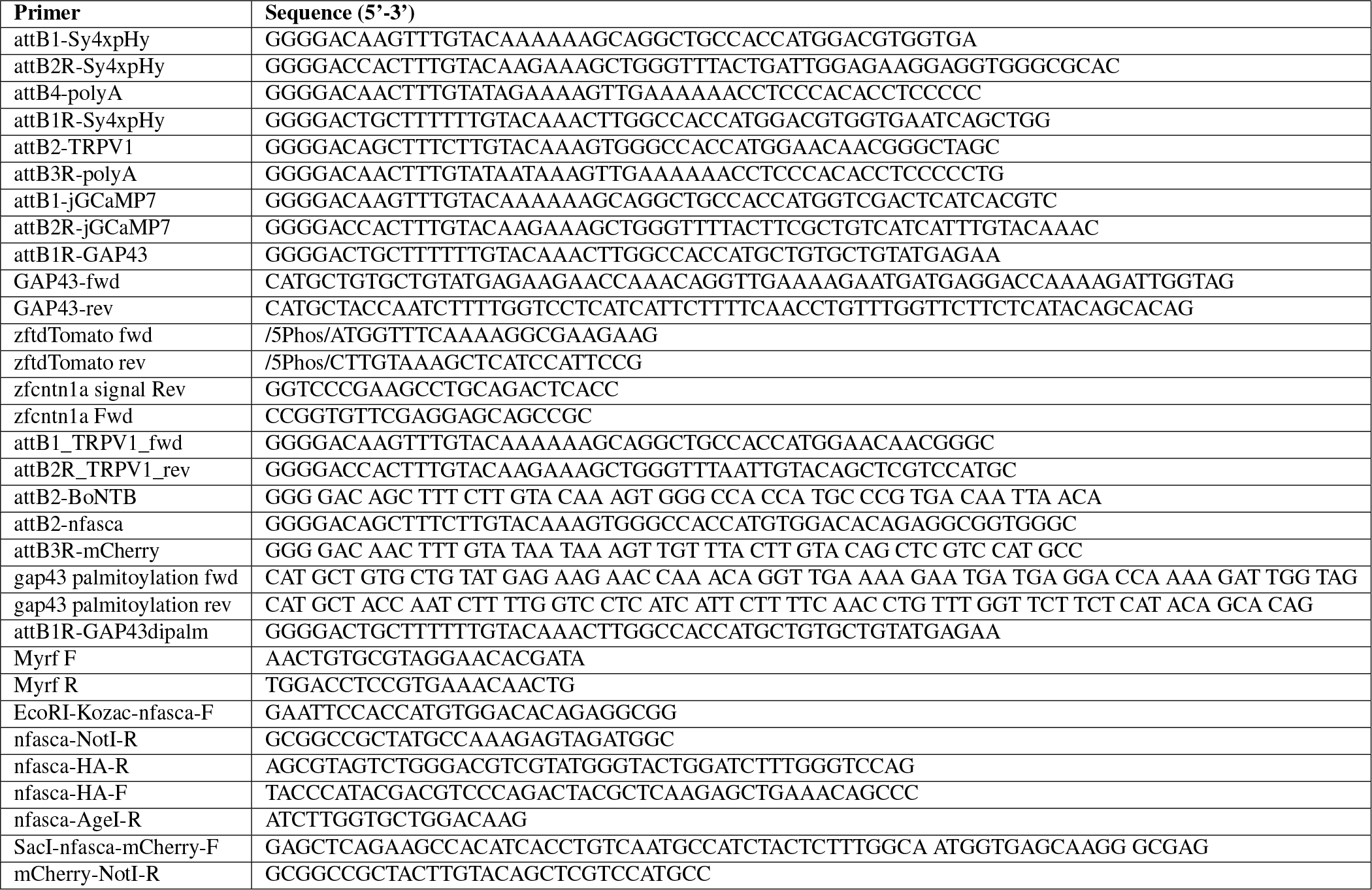
Primers

**Table S2 -.** BP reactions

| Entry vector | Template DNA | Fwd Primer | Rev Primer |
| --- | --- | --- | --- |
| pME-Sy4xpHy aka pME-SypHy | pcDNA3-SypHluorin 4x (22) (Addgene #37005) | attB1-Sy4xpHy | attB2R-Sy4xpHy |
| p5E-Sy4xpHy | UAS:Sy4xpHy (this study) | attB4-polyA | attB1R-Sy4xpHy |
| p3E-BoNTBmCherry | UAS:BoNTB-mCherry (29) | attB2-BoNTB | attB3R-polyA |
| p3E-nfasca-mCherry | pBH-UAS-HA-NF186-mCherry (this study) | attB2-nfasca | attB3R-polyA |
| p3E-TrpV1RFP | UAS:TrpV1-tagRFPt (this study) | attB2-TRPV1 | attB3R-polyA |
| pME-TrpV1-RFP | islet1:GAL4VP16,4xUAS:TRPV1-RFPt (52) | attB1_TRPV1_fwd | attB2_TRPV1_rev |
| pME-jGCaMP7s | pGP-CMV-jGCaMP7s (53) (Addgene #104463) | attB1-jGCaMP7 | attB2R-jGCaMP7 |
| p5E-axonjGCaMP7s | pME-axon-jGCaMP7s (this study) | attB4-polyA | attB1R-GAP43 |

**Table S3 -.**
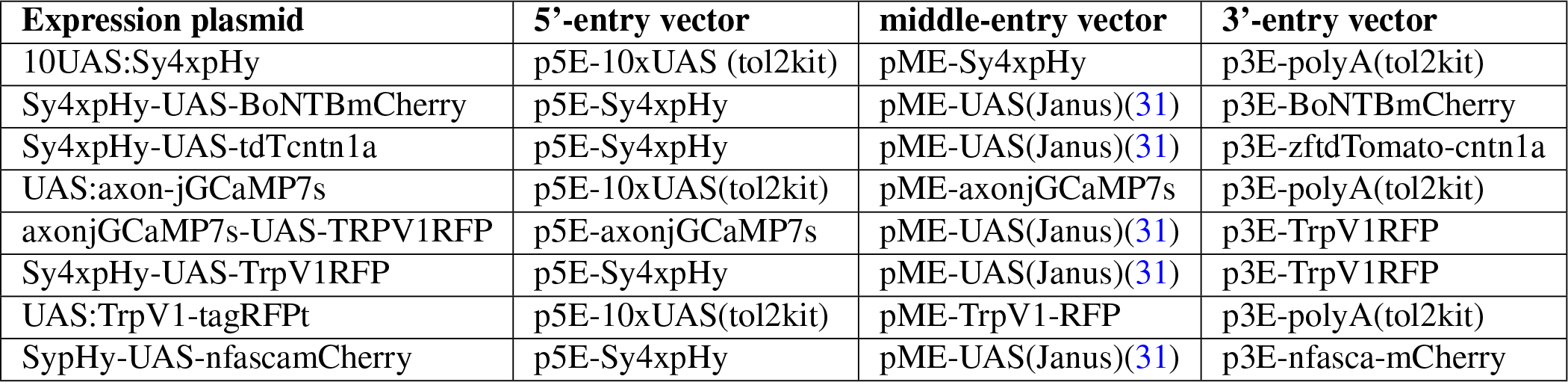
LR reactions

## Supplementary Figures

**Fig. S1.**
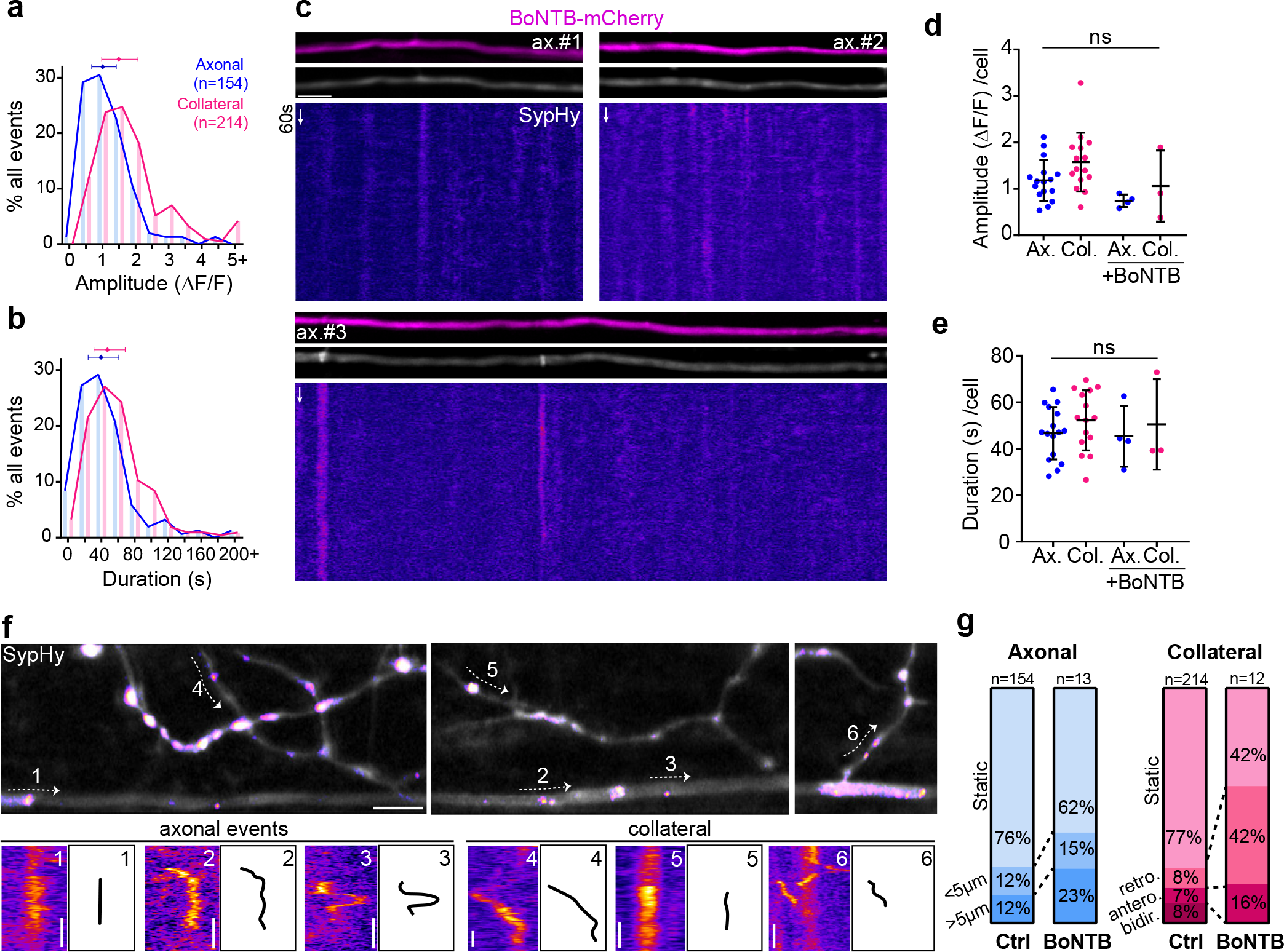
SypHy characterization and validation with BoNTB-mCherry, Related to Figure 1. a-b. Amplitude and duration distributions of axonal and collateral Syphy events largely overlap. Indicated number of events from 17 control axons from 16 animals; 15 BoNTB axons from 15 animals. c. Three examples of reticulospinal axons co-expressing SypHy and BoNTB-mCherry, which silences SypHy activity. d-e. SypHy event amplitude and duration per cell; note trend to amplitude reduction in remaining events in BoNTB+ axons. (d: p=0.05 Ax vs Col; p=0.07 Ax vs Ax+BoNTB; p=0.23 Col vs Col+BoNTB; Student’s t-test. e. p=0.21 Ax vs Col; p=0.84 Ax vs Ax+BoNTB; p=0.85 Col vs Col+BoNTB; Student’s t-test; 17 control axons from 16 animals; 15 BoNTB axons from 15 animal, averaged per animal). f-g. Kymographs of static and slightly displaced axonal and collateral events and quantification (binned into <5μm or >5μm cumulative displacement, retrograde, anterograde or bidirectional). Note proportional increase of dynamic events in silenced axons, including retrograde collateral and large displacement axonal events, suggesting they represent processes other than exocytosis (e.g. recycling endocytosis, longer-range transport). Scale bars: 5μm (c, f), 60s (c, f). Graphs display mean and standard deviation.

**Fig. S2.**
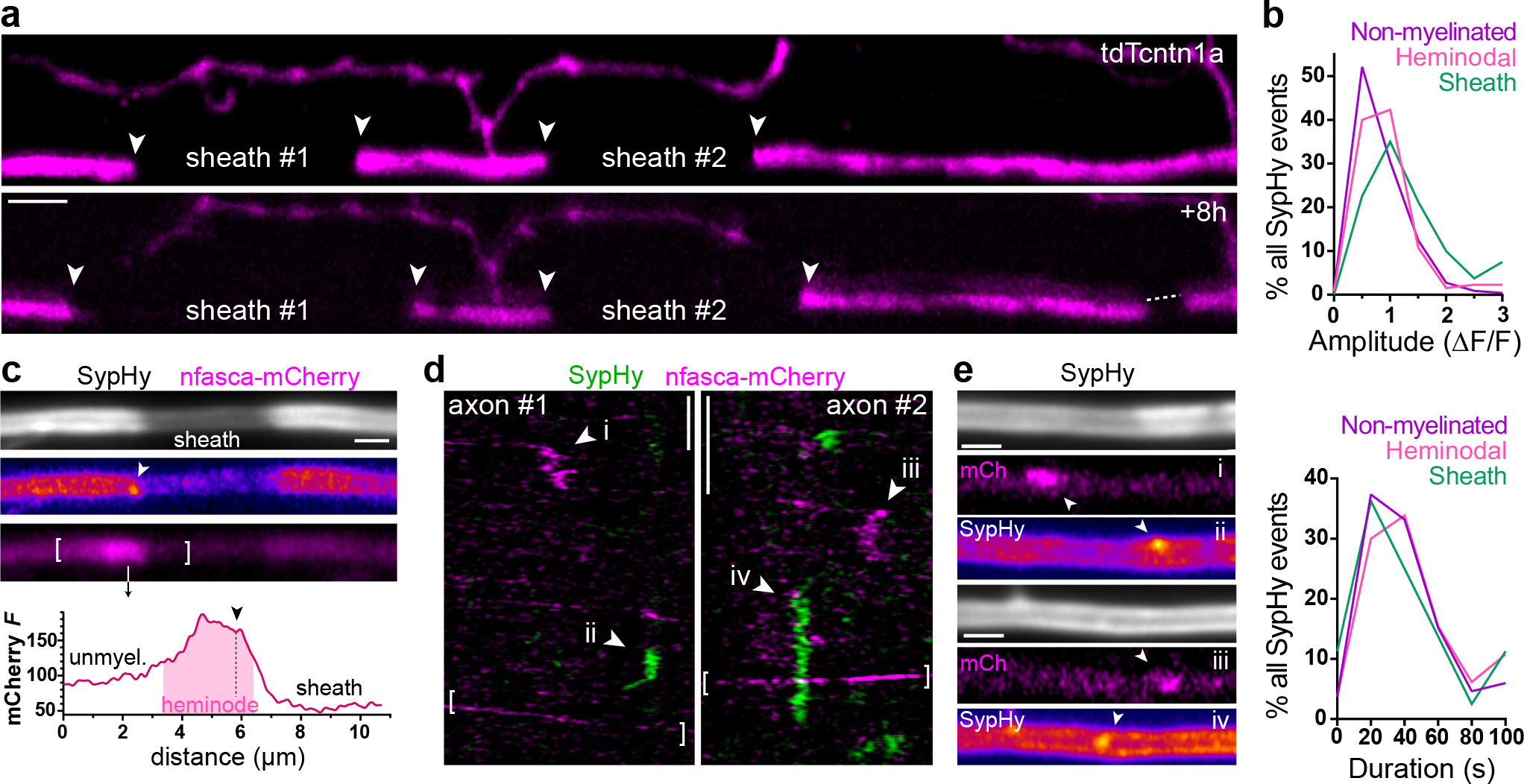
Axonal hotspots of SypHy at heminodal locations, Related to Figure 3. a. tdTcntn1a reveals sheaths and non-myelinated regions which can be followed over time. Panel below, 8 hours later, note sheath growth (arrowheads) and new sheath (dashed line). Related to Fig. 1 and 3. b. SypHy event amplitudes and durations are similar in all axonal locations (80 sheath events, 130 heminodal events, 217 non-myelinated events from 48 axons from 38 animals). c. nfasca-mCherry is enriched at putative heminodes (arrowhead, example heminodal SypHy event). Graph indicates mCherry fluorescence intensity profile (in AU) along bracketed region. d. Kymographs of axons co-expressing SypHy and nfasca-mCherry reveal independent transport, with distinct events labelled i-iv expanded in e. Brackets indicate transport particles. e. Frames from individual events indicated in d. Scale bars: 5μm (a), 2μm (c, e), 60s (d).

**Fig. S3.**
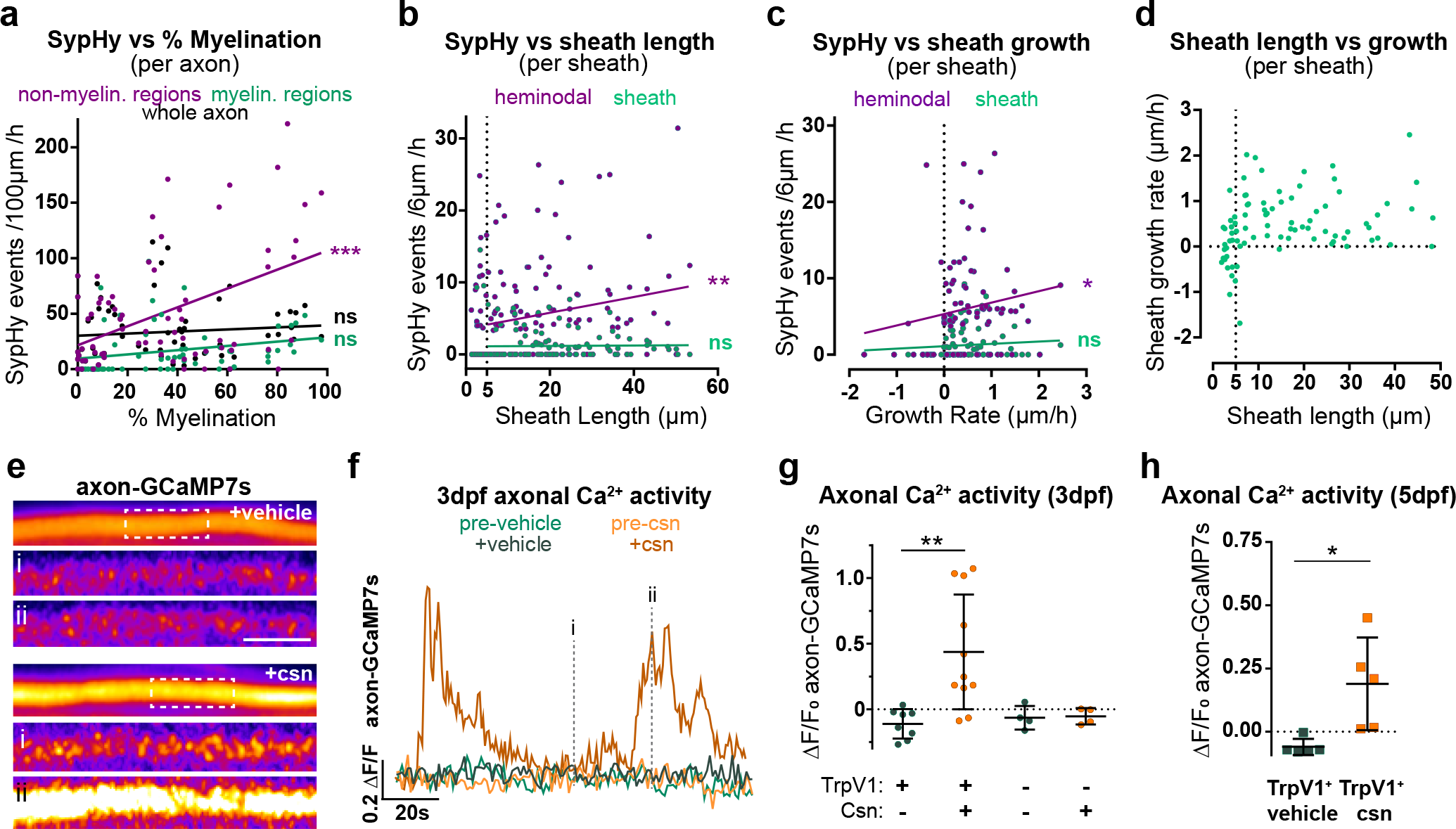
Temporal and causal relation of SypHy and sheath growth, Related to Figure 3. a. %myelination specifically correlates with SypHy frequency in nonmyelinated regions (Pearson r: 0.46, p<0.0001) but not in myelinated regions (Pearson r:0.25, p=0.05) or overall axonal SypHy (Pearson r:0.10, p=0.43). 70 myelinated axons from 57 animals. b. Sheath length positively correlates with SypHy frequency in heminodes (Pearson r: 0.23, p=0.003), but not under the sheath (Pearson r: 0.07, p=0.39). 162 sheaths from 61 axons from 54 animals. c. Monotonic correlation between sheath growth rate and SypHy frequency in heminodes (Spearman’s r:0.25, p=0.02) but not under the sheath (Spearman’s r:0.14, p=0.21). Pearson’s test of linear correlation ns; line drawn for illustration of correlation. 86 sheaths from 39 axons from 36 animals. d. Relation between sheath length and growth rate, note nascent <5μm sheaths may grow or shrink, but >5μm sheaths mostly grow (86 sheaths from 39 axons in 35 fish). e. Axon-GCaMP7s in 3dpf TrpV1 + axons, top panels are average projection, panels below are individual frames. Only csn increases Ca^2+^ activity. f. Fluorescence time-course of boxed ROIs in e. g-h. Quantification of average axonGCaMP7s activity at 3dpf (g) and 5dpf (h), showing specific increase in csn-treated TrpV1+ axons (3dpf: csn vs vehicle treatment of TrpV1+ axons: p=0.003, Student’s t-test, 19 TrpV1+ axons in 19 animals, 8 TrpV1-axons in 8 animals; 5dpf: p=0.017, 10 TrpV1+ axons in 10 animals). Scale bars: 2μm (e). Graphs display mean and standard deviation.

## Supplementary Movie Legends

**Movie S1 – SypHy activity along an individual reticulospinal axon (related to Fig. 1).** Raw fluorescence (top) and ΔF/F (bottom) in a SypHy-expressing reticulospinal axon in a wildtype larva at 4 days postfertilization, when myelination (not shown here) is ongoing. Arrowheads point to examples of SypHy events occurring along collaterals (where presynaptic terminals are located) and the main axonal projection. Time indicated in min:sec.

**Movie S2 – SypHy activity is silenced with BoNT-B co-expression (related to Fig. S1).** Raw fluorescence (top) and ΔF/F (bottom) in a single reticulospinal axon co-expressing SypHy and BoNTB-mCherry, showing essentially complete abrogation of axonal and collateral SypHy events. Time indicated in min:sec.

**Movie S3 – SypHy events are mostly static (related to Fig. S1).** Raw fluorescence (top) and ΔF/F (bottom) of three further examples of wildtype larva in which individual axons express SypHy. Note that SypHy events can be static (white arrowheads) or show displacement after their appearance (yellow arrowheads), quantified in Fig. S1. Time indicated in min:sec.

**Movie S4 – Relationship between SypHy activity and myelination (related to Fig. 1).** An individual reticulospinal axon co-expressing the tdTomato-cntn1a reporter of myelination, whose gaps indicate myelin sheaths, and SypHy reporter, enabling accurate quantification of SypHy activity along myelinated and unmyelinated regions. Arrowheads point to examples of SypHy events occurring along collaterals and axon. Time indicated in min:sec.

**Movie S5 - SypHy activity along an individual reticulospinal axon (related to Fig. 2) in a wildtype larva.** This larva is an homozygous wildtype sibling of a *myrf*^UE70^ litter. Time indicated in min:sec.

**Movie S6 - SypHy activity along an individual reticulospinal axon (related to Fig. 2) in an hypomyelinated myrf mutant larva.** This larva is an homozygous mutant sibling of a myrf^UE70^ litter. Note reduction in axonal, but not collateral SypHy events compared to wildtype sibling in Movie S5. Time indicated in min:sec.

**Movie S7 – Ca^2+^ response to capsaicin (csn) stimulation in a TrpV1+ expressing axon (related to Fig. S3).** Raw fluorescence of an individual reticulospinal axon co-expressing TrpV1-tagRFP and axon-GCaMP7s before (pre-csn) and during (+csn) capsaicin treatment. Note increase in axonal calcium activity. Time indicated in min:sec.

**Movie S8 – SypHy response to capsaicin (csn) stimulation in a TrpV1+ expressing axon (related to Fig. 3).** Raw fluorescence (top) and ΔF/F (bottom) of an individual reticulospinal axon co-expressing TrpV1-RFP and SypHy before (pre-csn) and during (+csn) capsaicin treatment. Note increase in SypHy activity. Time indicated in min:sec.

## Bibliography

1. Huxley, A. F & Stämpfli, R. Evidence for saltatory conduction in peripheral myelinated nerve fibres. The Journal of Physiology 108, 315–339 (1949). URL http://doi.wiley.com/10.1113/jphysiol.1949.sp004335.

2. Fünfschilling, U. et al. Glycolytic oligodendrocytes maintain myelin and long-term axonal integrity. Nature 485, 517–521 (2012). URL http://dx.doi.org/10.1038/nature11007.

3. Lee, Y. et al. Oligodendroglia metabolically support axons and contribute to neurodegeneration. Nature 487, 443–448 (2012). URL http://dx.doi.org/10.1038/nature11314.

4. Mukherjee, C. et al. Oligodendrocytes provide antioxidant defense function for neurons by secreting ferritin heavy chain. Cell Metabolism 32, 259–272.e10 (2020). URL http://www.sciencedirect.com/science/article/pii/S1550413120303004.

5. Almeida, R. G. The rules of attraction in central nervous system myelination. Frontiers in Cellular Neuroscience 12, 367 (2018). URL https://www.frontiersin.org/article/10.3389/fncel.2018.00367/full.

6. Williamson, J. M. & Lyons, D. A. Myelin dynamics throughout life: An ever-changing landscape? Frontiers in Cellular Neuroscience 12, 424 (2018). URL https://www.frontiersin.org/article/10.3389/fncel.2018.00424/full.

7. Bacmeister, C. M. et al. Motor learning promotes remyelination via new and surviving oligodendrocytes. BioRxiv (2020). URL http://biorxiv.org/lookup/doi/10.1101/2020.01.28.923656.

8. McKenzie, I. A. et al. Motor skill learning requires active central myelination. Science 346, 318–322 (2014). URL http://dx.doi.org/10.1126/science.1254960.

9. Liu, J. et al. Impaired adult myelination in the prefrontal cortex of socially isolated mice. Nature Neuroscience 15, 1621–1623 (2012). URL http://dx.doi.org/10.1038/nn.3263.

10. Makinodan, M., Rosen, K. M., Ito, S. & Corfas, G. A critical period for social experiencedependent oligodendrocyte maturation and myelination. Science 337, 1357–1360 (2012). URL http://dx.doi.org/10.1126/science.1220845.

11. Swire, M., Kotelevtsev, Y., Webb, D. J., Lyons, D. A. & Ffrench-Constant, C. Endothelin signalling mediates experience-dependent myelination in the CNS. eLife 8 (2019). URL http://dx.doi.org/10.7554/{eLife}.49493.

12. Pan, S., Mayoral, S. R., Choi, H. S., Chan, J. R. & Kheirbek, M. A. Preservation of a remote fear memory requires new myelin formation. Nature Neuroscience 23, 487–499 (2020). URL http://www.nature.com/articles/s41593-019-0582-1.

13. Steadman, P. E. et al. Disruption of oligodendrogenesis impairs memory consolidation in adult mice. Neuron 105, 150–164.e6 (2020). URL https://linkinghub.elsevier.com/retrieve/pii/S0896627319308864.

14. Wang, F. et al. Myelin degeneration and diminished myelin renewal contribute to age-related deficits in memory. Nature Neuroscience 23, 481–486 (2020). URL http://www.nature.com/articles/s41593-020-0588-8.

15. Almeida, R. G. & Lyons, D. A. On myelinated axon plasticity and neuronal circuit formation and function. The Journal of Neuroscience 37, 10023–10034 (2017). URL http://www.jneurosci.org/lookup/doi/10.1523/{JNEUROSCI}.3185-16.2017.

16. Hines, J. H., Ravanelli, A. M., Schwindt, R., Scott, E. K. & Appel, B. Neuronal activity biases axon selection for myelination in vivo. Nature Neuroscience 18, 683–689 (2015). URL http://dx.doi.org/10.1038/nn.3992.

17. Koudelka, S. et al. Individual neuronal subtypes exhibit diversity in CNS myelination mediated by synaptic vesicle release. Current Biology 26, 1447–1455 (2016). URL http://dx.doi.org/10.1016/j.cub.2016.03.070.

18. Mensch, S. et al. Synaptic vesicle release regulates myelin sheath number of individual oligodendrocytes in vivo. Nature Neuroscience 18, 628–630 (2015). URL http://dx.doi.org/10.1038/nn.3991.

19. Hughes, A. N. & Appel, B. Oligodendrocytes express synaptic proteins that modulate myelin sheath formation. Nature Communications 10, 4125 (2019). URL http://www.nature.com/articles/s41467-019-12059-y.

20. Micu, I. et al. The molecular physiology of the axo-myelinic synapse. Experimental Neurology 276, 41–50 (2016). URL http://dx.doi.org/10.1016/j.expneurol.2015.10.006.

21. Marisca, R. et al. Functionally distinct subgroups of oligodendrocyte precursor cells integrate neural activity and execute myelin formation. Nature Neuroscience 23, 363–374 (2020). URL http://www.nature.com/articles/s41593-019-0581-2.

22. Zhu, Y., Xu, J. & Heinemann, S. F. Two pathways of synaptic vesicle retrieval revealed by single-vesicle imaging. Neuron 61, 397–411 (2009). URL http://dx.doi.org/10.1016/j.neuron.2008.12.024.

23. Jahn, R. & Südhof, T. C. Synaptic vesicles and exocytosis. Annual Review of Neuroscience 17, 219–246 (1994). URL http://dx.doi.org/10.1146/annurev.ne.17.030194.001251.

24. Südhof, T. C., Lottspeich, F., Greengard, P., Mehl, E. & Jahn, R. A synaptic vesicle protein with a novel cytoplasmic domain and four transmembrane regions. Science 238, 1142–1144 (1987). URL http://dx.doi.org/10.1126/science.3120313.

25. Wiedenmann, B. & Franke, W. W. Identification and localization of synaptophysin, an integral membrane glycoprotein of mr 38,000 characteristic of presynaptic vesicles. Cell 41, 1017–1028 (1985). URL http://dx.doi.org/10.1016/s0092-8674(85)80082-9.

26. Granseth, B., Odermatt, B., Royle, S. J. & Lagnado, L. Clathrin-mediated endocytosis is the dominant mechanism of vesicle retrieval at hippocampal synapses. Neuron 51, 773–786 (2006). URL http://dx.doi.org/10.1016/j.neuron.2006.08.029.

27. Pan, P.-Y., Marrs, J. & Ryan, T. A. Vesicular glutamate transporter 1 orchestrates recruitment of other synaptic vesicle cargo proteins during synaptic vesicle recycling. The Journal of BiologicalChemistry 290, 22593–22601 (2015). URL http://dx.doi.org/10.1074/jbc.M115.651711.

28. Gahtan, E. & O’Malley, D. M. Visually guided injection of identified reticulospinal neurons in zebrafish: a survey of spinal arborization patterns. The Journal of Comparative Neurology 459, 186–200 (2003). URL http://dx.doi.org/10.1002/cne.10621.

29. Sternberg, J. R. et al. Optimization of a neurotoxin to investigate the contribution of excitatory interneurons to speed modulation in vivo. Current Biology 26, 2319–2328 (2016). URL http://dx.doi.org/10.1016/j.cub.2016.06.037.

30. Schiavo, G. et al. Tetanus and botulinum-b neurotoxins block neurotransmitter release by proteolytic cleavage of synaptobrevin. Nature 359, 832–835 (1992). URL http://dx.doi.org/10.1038/359832a0.

31. Almeida, R. G. et al. Myelination of neuronal cell bodies when myelin supply exceeds axonal demand. Current Biology 28, 1296–1305.e5 (2018). URL http://linkinghub.elsevier.com/retrieve/pii/S0960982218302549.

32. Salzer, J. L. Polarized domains of myelinated axons. Neuron 40, 297–318 (2003). URL http://dx.doi.org/10.1016/s0896-6273(03)00628-7.

33. Zhang, Y. et al. Assembly and maintenance of nodes of ranvier rely on distinct sources of proteins and targeting mechanisms. Neuron 73, 92–107 (2012). URL http://dx.doi.org/10.1016/j.neuron.2011.10.016.

34. Zonta, B. et al. Glial and neuronal isoforms of neurofascin have distinct roles in the assembly of nodes of ranvier in the central nervous system. The Journal of Cell Biology 181, 1169–1177 (2008). URL http://dx.doi.org/10.1083/jcb.200712154.

35. Broussard, G. J. et al. In vivo measurement of afferent activity with axon-specific calcium imaging. Nature Neuroscience 21, 1272–1280 (2018). URL http://www.nature.com/articles/s41593-018-0211-4.

36. Chang, K.-J., Redmond, S. A. & Chan, J. R. Remodeling myelination: implications for mechanisms of neural plasticity. Nature Neuroscience 19, 190–197 (2016). URL http://dx.doi.org/10.1038/nn.4200.

37. Mayoral, S. R. & Chan, J. R. The environment rules: spatiotemporal regulation of oligodendrocyte differentiation. Current Opinion in Neurobiology 39, 47–52 (2016). URL http://dx.doi.org/10.1016/j.conb.2016.04.002.

38. Normand, E. A. & Rasband, M. N. Subcellular patterning: axonal domains with specialized structure and function. Developmental Cell 32, 459–468 (2015). URL http://dx.doi.org/10.1016/j.devcel.2015.01.017.

39. Susuki, K. et al. Three mechanisms assemble central nervous system nodes of ranvier. Neuron 78, 469–482 (2013). URL http://dx.doi.org/10.1016/j.neuron.2013.03.005.

40. Bekku, Y. & Salzer, J. L. Independent anterograde transport and retrograde cotransport of domain components of myelinated axons. The Journal of Cell Biology 219 (2020). URL http://dx.doi.org/10.1083/jcb.201906071.

41. Gibson, E. M. et al. Neuronal activity promotes oligodendrogenesis and adaptive myelination in the mammalian brain. Science 344, 1252304 (2014). URL http://dx.doi.org/10.1126/science.1252304.

42. Mitew, S. et al. Pharmacogenetic stimulation of neuronal activity increases myelination in an axon-specific manner. Nature Communications 9, 306 (2018). URL http://dx.doi.org/10.1038/s41467-017-02719-2.

43. Brownstone, R. M. & Chopek, J. W. Reticulospinal systems for tuning motor commands. Frontiers in Neural Circuits 12, 30 (2018). URL http://dx.doi.org/10.3389/fncir.2018.00030.

44. Jordan, L. M., Liu, J., Hedlund, P. B., Akay, T. & Pearson, K. G. Descending command systems for the initiation of locomotion in mammals. Brain Research Reviews 57, 183–191 (2008). URL http://dx.doi.org/10.1016/j.brainresrev.2007.07.019.

45. Lundgaard, I. et al. Neuregulin and BDNF induce a switch to NMDA receptor-dependent myelination by oligodendrocytes. PLoS Biology 11, e1001743 (2013). URL http://dx.doi.org/10.1371/journal.pbio.1001743.

46. Nusslein-Volhard, C. & Dahm, R. Zebrafish (Oxford University Press, 2002).

47. Westerfield, M. The zebrafish book: a guide for the laboratory use of zebrafish. http://zfin.org/zf_info/zfbook/zfbk.html (2000). URL https://ci.nii.ac.jp/naid/10029409142/.

48. Kimmel, C. B., Ballard, W. W., Kimmel, S. R., Ullmann, B. & Schilling, T. F. Stages of embryonic development of the zebrafish. Developmental Dynamics 203, 253–310 (1995). URL http://dx.doi.org/10.1002/aja.1002030302.

49. Antinucci, P., Folgueira, M. & Bianco, I. H. Pretectal neurons control hunting behaviour. eLife 8 (2019). URL http://dx.doi.org/10.7554/{eLife}.48114.

50. Kwan, K. M. et al. The tol2kit: a multisite gateway-based construction kit for tol2 transposon transgenesis constructs. Developmental Dynamics 236, 3088–3099 (2007). URL http://dx.doi.org/10.1002/dvdy.21343.

51. Distel, M., Hocking, J. C., Volkmann, K. & Köster, R. W. The centrosome neither persistently leads migration nor determines the site of axonogenesis in migrating neurons in vivo. The Journal of Cell Biology 191, 875–890 (2010). URL http://dx.doi.org/10.1083/jcb.201004154.

52. Chen, S., Chiu, C. N., McArthur, K. L., Fetcho, J. R. & Prober, D. A. TRP channel mediated neuronal activation and ablation in freely behaving zebrafish. Nature Methods 13, 147–150 (2016). URL http://dx.doi.org/10.1038/nmeth.3691.

53. Dana, H. et al. High-performance calcium sensors for imaging activity in neuronal populations and microcompartments. Nature Methods 16, 649–657 (2019). URL http://www.nature.com/articles/s41592-019-0435-6.

